# Microglial Calcium Signaling is Attuned to Neuronal Activity

**DOI:** 10.1101/2019.12.16.876060

**Authors:** Anthony D. Umpierre, Lauren L. Bystrom, Yanlu Ying, Yong U. Liu, Long-Jun Wu

## Abstract

Microglial calcium signaling underlies a number of key physiological processes *in situ*, but has not been studied *in vivo* in an awake animal where neuronal function is preserved. Using multiple GCaMP6 variants targeted to microglia, we assessed how microglial calcium signaling responds to alterations in neuronal activity across a wide physiological range. We find that only a small subset of microglial somata and processes exhibited spontaneous calcium transients. However, hyperactive and hypoactive shifts in neuronal activity trigger increased microglial process calcium signaling, often concomitant with process extension. On the other hand, changes in somatic calcium activity are only observed days after severe seizures. Our work reveals that microglia have highly distinct microdomain signaling, and that processes specifically respond to bi-directional shifts in neuronal activity through calcium signaling.

## INTRODUCTION

Microglia are resident immune cells of the CNS, which are specialized to respond to both immunological and neuronal stimuli. Early work *in situ* suggested that calcium activity was critical for microglia, with calcium signaling underlying key physiological functions such as motility, cytokine release, and receptor trafficking/diffusion (Färber and Kettenmann, 2006; Korvers et al., 2016; Toulme and Khakh, 2012). However, more recent *in vivo* studies have demonstrated that spontaneous calcium signaling is nearly absent in microglia (Brawek et al., 2017; Eichhoff et al., 2011; Pozner et al., 2015). Such observations downplay the potential utility of calcium signaling for microglial function.

Over the past two decades of astrocyte calcium research, it is become clear that microdomains (soma, major branches, and minor branches) have exquisitely different calcium signaling properties, with the smallest sub-compartments of the cell having the most frequent calcium signaling (Srinivasan et al., 2015). Microglial processes are well known to dynamically extend and retract as they survey the brain microenvironment (Nimmerjahn et al., 2005; Wake et al., 2009). To date, microglial processes have not been extensively evaluated for their calcium activity *in vivo*, nor has any assessment been performed in the awake animal. Herein, we evaluated microglial soma and process calcium signaling in the awake animal using 2-photon microscopy.

One of the key questions in microglial research is the extent to which microglia functionally respond to neuronal activity in the mature brain. Testing a high signal-to-noise GCaMP variant (GCaMP6s) and a membrane-tethered GCaMP variant (Lck-GCaMP6f), we find that microglia have very infrequent calcium activity in their processes under basal conditions. However, we find that microglial process calcium activity is strongly attuned to neuronal activity changes, with both neuronal hypoactivity and hyperactivity triggering increases in microglial process calcium signals. By contrast, microglial somatic calcium changes may represent a disease signature as somatic calcium changes are only evident in longitudinal epilepsy development. Our work reveals that microglia have highly dynamic process calcium activity during network activity shifts, most closely associated with processes undergoing extension.

## RESULTS

### Spontaneous microglial calcium activity reported in the Lck-GCaMP6f and cytosolic GCaMP6s mouse

Microglial calcium activity has been frequently described *in situ*, with limited studies performed *in vivo*. We first assessed microglial calcium activity in acute window preparations (24 h after surgery), testing the utility of two sensors: GCaMP6s (highest signal-to-noise ratio), and Lck-GCaMP6f (membrane tethered variant with idealized localization to processes) (Madisen et al., 2015; Srinivasan et al., 2016). Spontaneous calcium activity was studied in the awake, head-restrained animal (Figure 1A), using two-photon microscopy to assess microglia in layer I and II/III of somatosensory cortex. Microglial calcium activity was present in both the Lck-GCaMP6f mouse and the cytosolic GCaMP6s mouse (Figure 1B and 1C). However, the higher fluorescence amplitude and slower decay kinetics in the GCaMP6s animal made this fluorophore ideal for studying calcium activity in thin microglial processes. On average, GCaMP6s reported calcium transients with a 50% greater amplitude and a 60% increase in signal area (Figure 1D). For these reasons, we could detect calcium events in a greater percentage of microdomains in the GCaMP6s animal (Figure 1C). Therefore, we adopted GCaMP6s mice for our studies of microglial calcium activity. It should be mentioned, however, that Lck-GCaMP6f mice do report unique phenomenon not observed in the GCaMP6s mouse. Specifically, small amplitude, widespread, and temporally coordinated calcium events can be observed in the parenchyma of Lck-GCaMP6f mice using a grid-based analysis (Figure 1E).

**Figure 1:**
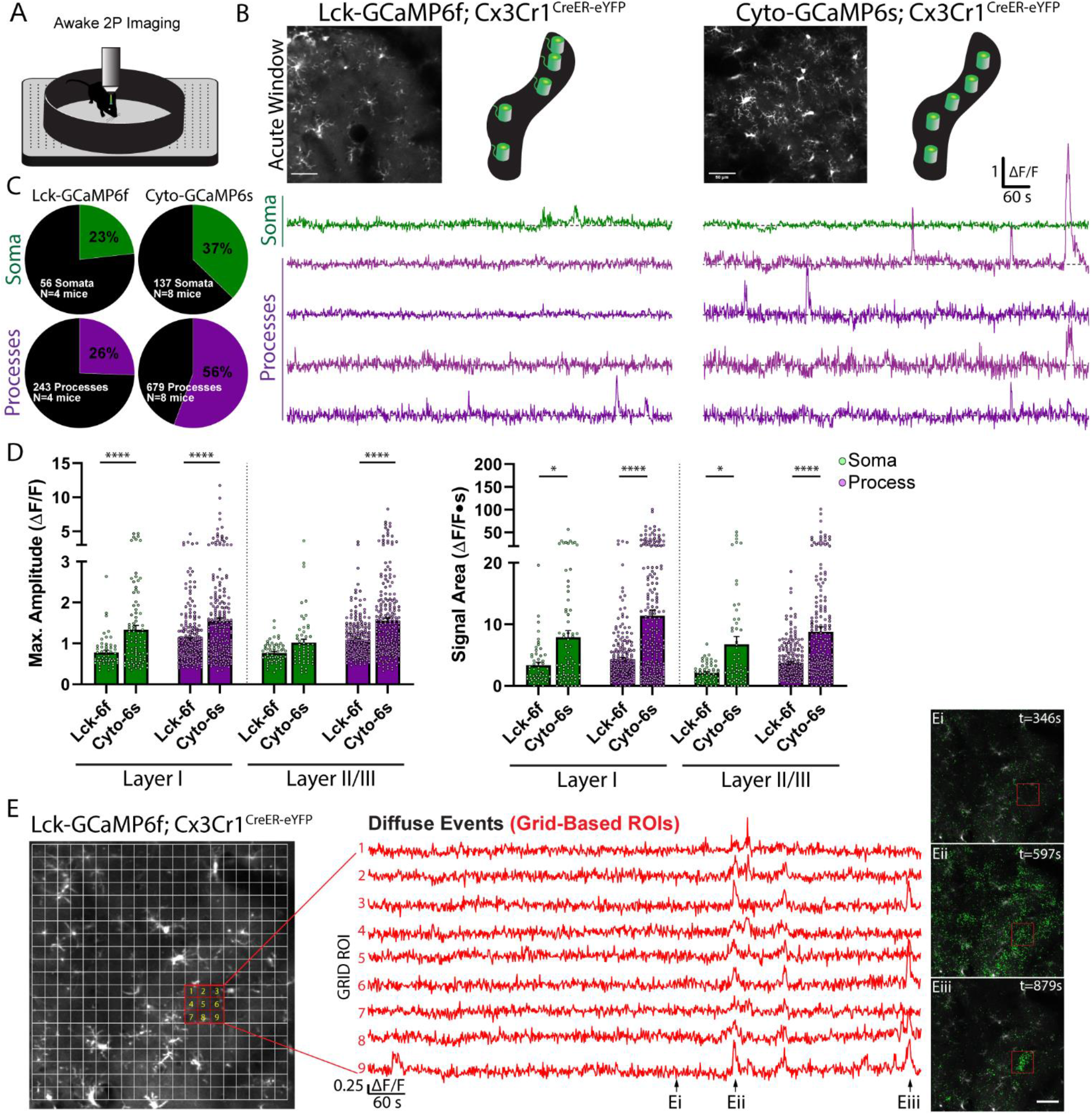
Spontaneous microglial calcium activity reported in the Lck-GCaMP6f and cytosolic GCaMP6s mouse. **(A)** Imaging setup: *in vivo* two-photon imaging of microglial calcium activity in the awake, head-restrained animal. **(B)** Average intensity projection images and microglial ΔF/F calcium traces reported by Lck-GCaMP6f or cytosolic-GCaMP6s mice. **(C)** Percentage of active microdomains (STAR Methods) reported by each sensor. **(D)** Maximum amplitude and signal area reported by each sensor (bar: mean ± SEM; dots: individual microdomains; 1-Way ANOVA with Sidak’s post-hoc comparison). **(E)** The Lck-GCaMP6f mouse reports microglial calcium activity that is widespread, coordinated, and low-level throughout the parenchyma (Eii), or concentrated within a smaller region (Eiii). Scale bar: 50 μm (B and E). *p<0.05, ****p<0.0001. N=4 Lck-GCaMP6f mice, N=8 Cyto-GCaMP6s mice. See also Figure S1.

Notably, as immune cells, microglia respond to a variety of stimuli, including injury and inflammation. We next assessed whether studying microglial calcium activity soon after surgery in the acute window preparation (<24 hr) resulted in similar calcium activity patterns to a chronic window animal (4 weeks after surgery; Figure S1A). Notably, the morphology of cortical microglia was highly similar in acute and chronic window studies (Sholl analysis, Figure S1B). Despite the similarities in morphology, microglial process calcium activity was significantly greater in acute window preparations, while somatic calcium activity levels were indistinguishable (Figure S1C-S1G). Approximately one-third of microglia somata exhibited calcium activity in layer I (55-75 μm depth) and layer II/III (150-170 μm depth) in both the acute and chronic window preparation (Figure S1E). However, the proportion of active processes dropped for both layer I microglia (56% to 35%), and layer II/III microglia (54% to 41%, Figure S1E). In the acute window preparation, a subset of processes with high calcium activity was present across both cortical depths; however, this population was not maintained one month after surgery (Figure S1G, top axis of signal area and cumulative distribution curves). Notably, in the chronic window animal, baseline recordings from the same region were stable over three successive days of recording (Figure S1H). For these reasons, we performed all other studies in chronic window mice to avoid any potential confounds associated with post-surgery microglial activation (Xu et al., 2007).

### Isoflurane anesthesia increases microglial process outgrowth and calcium activity

To date, it is unclear whether neuronal activity influences microglial calcium signaling. We first assessed whether general anesthesia impacts microglial calcium signaling. After recording baseline calcium activity in the awake animal, we introduced isoflurane anesthesia via nose cone (Figure 2A). Importantly, the isoflurane concentration chosen for maintenance (1.5-2% in oxygen) is intended to mimic strong sensory blockade during surgery and is higher than the concentration typically used during anesthetized imaging (0.5-1.5%). Our studies of neuronal activity indicate that 1.5-2% isoflurane maintenance can reduce excitatory calcium signaling by 91 ± 0.11% in the somatosensory cortex (AAV.CaMKIIa.GCaMPs.WPRE transfected mice, layer II/III of somatosensory cortex; Figures 2B and 2C). In response to strongly reduced neuronal activity, microglia demonstrate an increase in process calcium activity (Figures 2D, 2E, and Video S1). Microglial process calcium activity reaches peak levels approximately 10 minutes after isoflurane induction. Thereafter, microglial processes display a 290 ± 29% increase in average ΔF/F calcium signal area (Figure 2F). Despite strong increases in process calcium activity, somatic calcium signaling remained unaffected in microglia under general anesthesia, with no changes in the proportion or signal area of somatic events (Figure 2F). These results provide early evidence that microglial compartments could have distinct calcium-associated functions.

**Figure 2:**
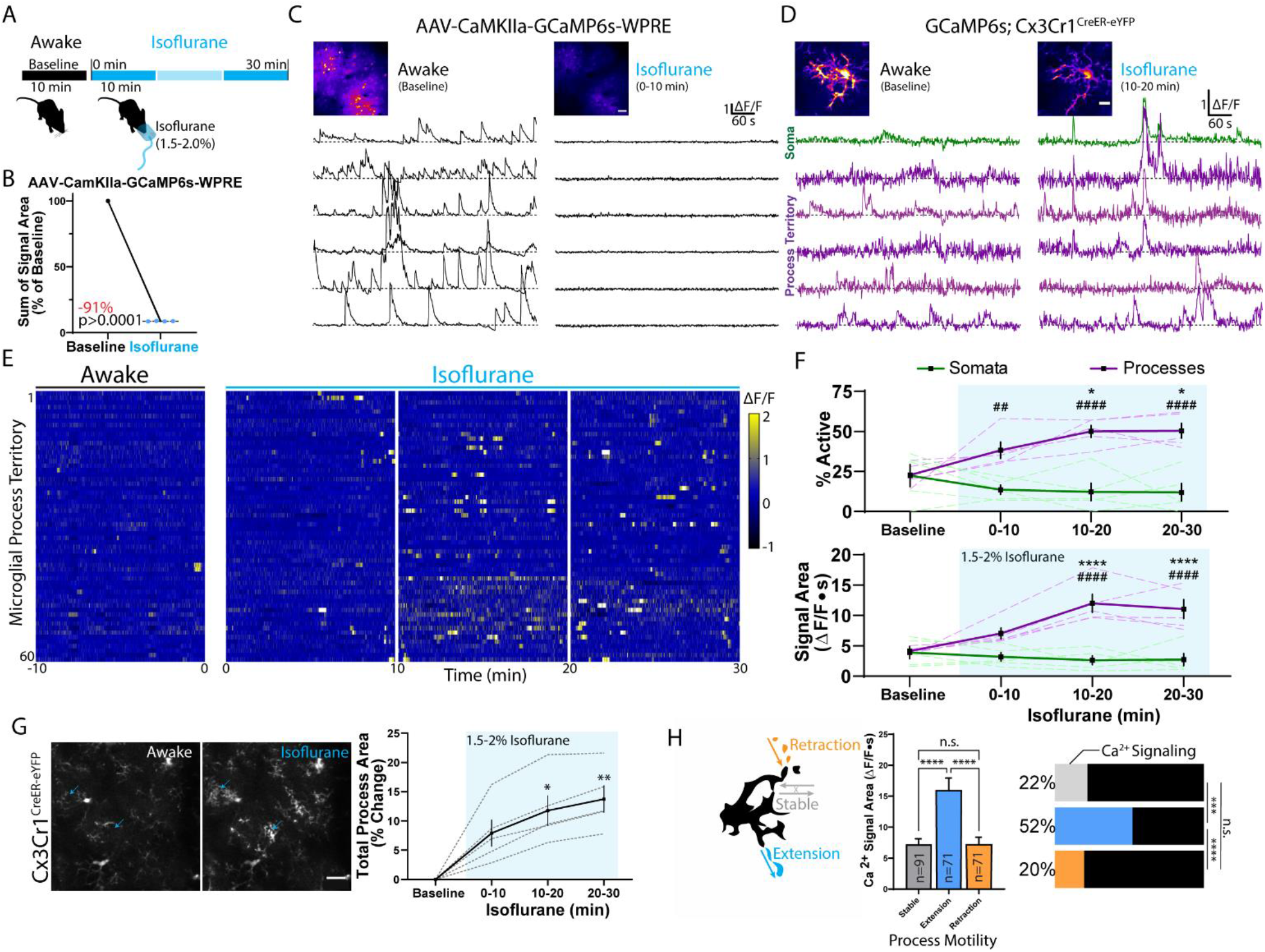
Isoflurane anesthesia increases microglial process outgrowth and calcium activity. **(A)** Outline of microglial calcium imaging before and during isoflurane. **(B-C)** Excitatory neuronal calcium activity before and after isoflurane induction, including (B) baseline-normalized calcium activity under isoflurane (Paired t-test; dots: one animal), and (C) representative ΔF/F traces. **(D-E)** Microglial calcium activity under awake and isoflurane conditions, including ΔF/F traces from a representative cell (D) and heat map data from a representative animal (E). **(F)** Microdomains active and their signal areas from baseline through anesthesia (2-Way ANOVA; *Dunnett’s post hoc vs. baseline; #somata vs. processes). **(G)** Cx3Cr1^CreER-eYFP^ morphology in the awake animal and during isoflurane anesthesia. Summary of process area changes (1-Way ANOVA with Dunnett’s post-hoc vs. baseline). **(H)** Microglial process calcium activity for processes undergoing extension, retraction, or remaining stable (1-Way ANOVA with Tukey’s post-hoc comparison). Percentage of processes with each motility behavior also having calcium activity (Fisher’s exact test). Scale bars: 50 μm (C), 10 μm (D), 20 μm (G). N=4 AAV-CaMKIIa-GCaMP6s mice (B, C), N=5 GCaMP6s;Cx3Cr1^CreER-eYFP^ mice (D-H). Grouped data represent mean ± SEM; dashed lines represent individual animals (F, G). Numbers of processes surveyed in (H) are shown in the bars. *p<0.05, ##p<0.01, ***p<0.001, ****, ####p<0.0001.

In accordance with our previous studies in the awake animal, general anesthesia increases microglial process extension (Liu et al., 2019). At the isoflurane concentration used, we observed a gradual increase in total process area co-occurring with increased process calcium activity (Figure 2G). *In situ*, microglial process motility has been strongly associated with calcium activity (Langfelder et al., 2015; Nolte et al., 1996). *In vivo*, we find that microglial processes undergoing extension have the largest calcium signals and the highest association with calcium activity, when compared to processes that undergo retraction or remain stable (Figure 2H). However, extension may not categorically require calcium activity, as it is associated with only half of all extending processes. These findings suggest that structural changes alone are only partially explanatory of calcium activity under general anesthesia.

### Kainate administration leads to acute and longitudinal modulation of microglial calcium signaling

We additionally used kainate, an excitatory receptor agonist and chemoconvulsant, to determine if strong increases in neuronal activity can alter microglial calcium signaling. In a first cohort of chronic window mice, we recorded microglial calcium activity at baseline and soon after kainate-induced generalized seizures in awake mice (Figure S2A). Seizure generalization involves activation of cortical neuronal populations. In neuronal studies (AAV.CaMKIIa.GCaMPs.WPRE transfected mice), kainate increased excitatory calcium activity both soon after administration (15-30 min) and after the first observed seizure (55-fold increase in signal area; Figures S2B and S2C). After the first observed seizure, microglial process calcium activity was increased by 360 ± 26%, with a greater proportion of processes exhibiting calcium activity (32% to 73%; Figures S2D and S2E). Despite strong increases in process calcium signaling, microglial somatic signaling did not increase significantly as measured through average signal area; however, there was an increase in the proportion of somata exhibiting calcium activity (29% to 44%; Figures S2D and S2E). During kainate-induced seizures, the predominant motility behavior was process extension, with process area increasing across the parenchyma (Figure S2F). These findings demonstrate that both an approach to rapidly decrease neuronal activity (isoflurane) and an approach to rapidly increase neuronal activity (kainate) similarly result in large-scale increases in microglial process calcium activity. In addition, both approaches resulted in similar structural changes (process extension) and were insufficient to alter microglial somatic calcium activity.

The prolonged seizure state induced by kainate, called status epilepticus, can also serve as a precipitating insult for the later acquisition of epilepsy (Puttachary et al., 2015; Tse et al., 2014; Umpierre et al., 2016). In a second cohort of chronic window mice, we recorded baseline microglial calcium activity and then monitored status epilepticus severity after kainate administration (Figure 3A). Mice with more severe status epilepticus (≥8 generalized seizures) were longitudinally studied over a 14-day period for changes in microglial calcium signaling. For up to 7 days after kainate status epilepticus, layer I microglia displayed an altered morphology, marked by greater process ramification (Figures 3B, S3A, and S3B). Over the same 7-day time course, a greater proportion of microglial somata and processes displayed calcium activity, with increased calcium signaling lasting up to 10 days (Figures 3C-3E). During this period, microglial somata and processes could sustain high amplitude (>2 fold ΔF/F) calcium transients for exceptionally long periods (1-4 minutes; Figure 3C). Calcium activity was also highly synchronized between the processes and soma, best exemplified by the appearance of large, spreading microglial calcium waves (Figure 3D and Video S2). For these reasons, it is not surprising that the proportion of active microdomains and their signal areas are highly similar during this 14-day period (Figure 3E). Both somata and processes showed large increases in calcium activity before returning to near-baseline levels two weeks later (Figure 3E). Similar to prior experiments, we find that microglial processes undergoing extension have highly increased calcium activity, compared to processes undergoing retraction or remaining stable (Figures 3F, 3G, and Video S3). Thus, during a state of network rearrangement and inflammation, microglia display profound increases in their calcium signaling in both somata and processes.

**Figure 3:**
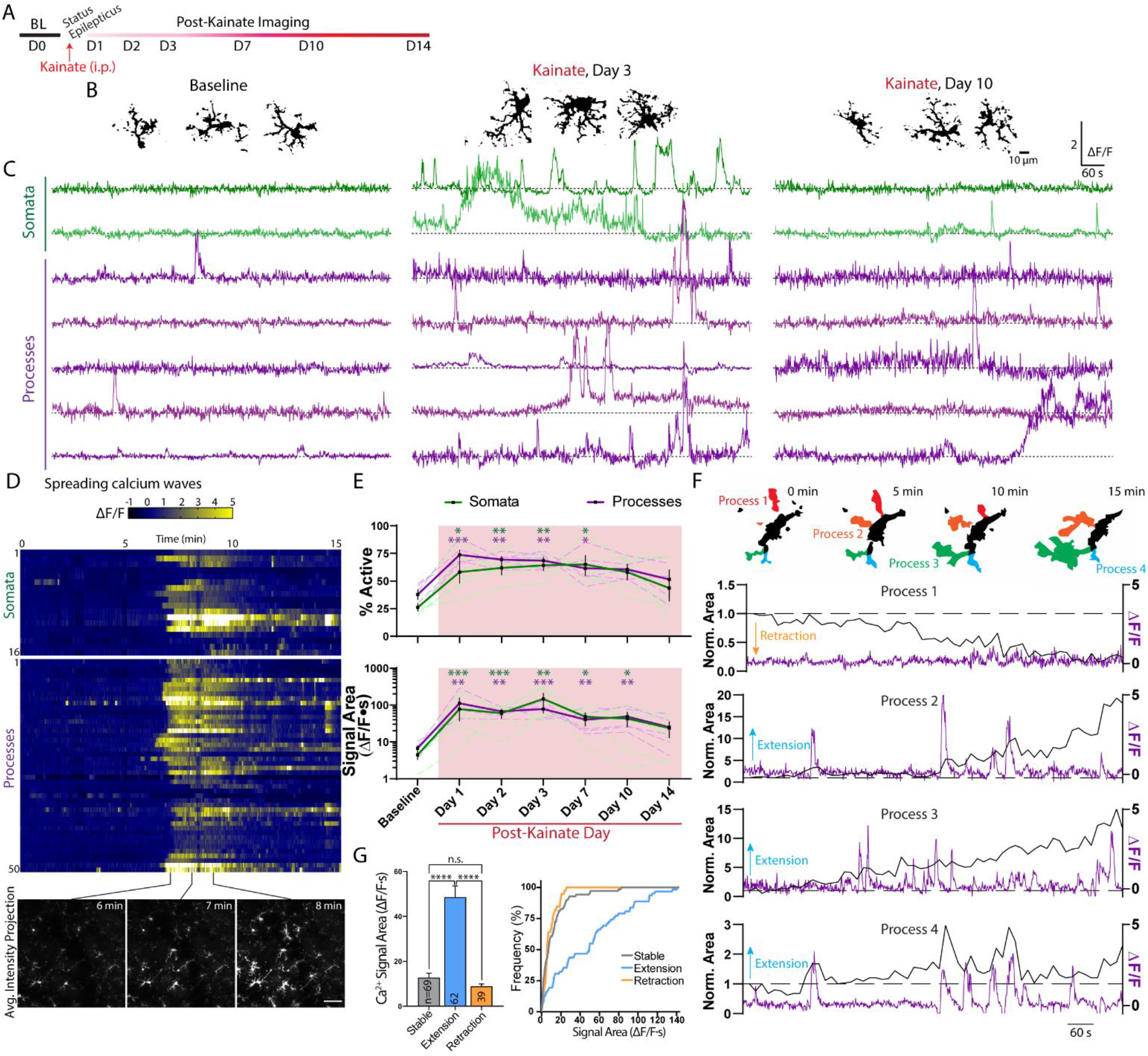
Kainate administration leads to longitudinal modulation of microglial calcium signaling. **(A)** Timeline of experiment. **(B)** Representative images of microglia morphology at baseline and following kainate status epilepticus. See also Figure S3. **(C)** ΔF/F traces of microglial somatic and process calcium activity. **(D)** Heat map of a microglial spreading calcium wave one day after kainate status epilepticus. See also Video S2. **(E)** Microdomains active and their signal areas (2-Way ANOVA with Dunnett’s post-hoc vs. baseline). **(F)** Representative changes in microglial process area correlated with ΔF/F calcium activity. See also Video S3. **(G)** Microglial process calcium activity for processes undergoing extension, retraction, or remaining stable (bar graph: mean ± SEM, 1-Way ANOVA with Tukey’s post-hoc testing; line graph: cumulative distribution curves). Scale bar: 50 μm (D). N=5 GCaMP6s;Cx3Cr1^CreER-eYFP^ mice. Grouped data represent mean ± SEM; dashed lines represent individual animals (E). Numbers of processes surveyed in (G) are shown in the bars. *p<0.05, **p<0.01, ***p<0.001. See also Figure S2.

### DREADD-based modulation of excitatory neuronal activity is sufficient to induce microglial calcium signaling

Isoflurane- and kainate-based changes in neuronal activity represent two of the more extreme shifts in network function. We additionally employed chemogenetic approaches to achieve smaller, more direct alterations to network function. DREADDs (Designer Receptors Exclusively Activated by Designer Drugs) utilize the expression of exogenous G-protein receptors to modulate cellular activity (Roth, 2016). We used a Gi system (hM4D) targeted to local CaMKIIa neurons of the somatosensory cortex to decrease neuronal activity and a Gq system (hM3D) to increase neuronal activity (Figure S4A). In a first series of experiments, we tested multiple doses of the DREADD ligand, Clozapine N-oxide (CNO), with the goal of finding a dose that could produce modest (30-40%) and intermediate (60-70%) changes in excitatory neuronal calcium activity. We also examined the time of peak effect in both systems between 15-60 min after injection (Figure S4A, timeline). In the Gi system, a 2.5 mg/kg (i.p.) dose of CNO decreased CaMKIIa calcium signaling by 51 ± 4.2%, while a 5.0 mg/kg dose decreased signaling by 66 ± 3.5%. Due to the similar responses, a 1.25 mg/kg (i.p.) dose of CNO was also tested, resulting in a 31 ± 2.7% decrease in CaMKIIa calcium activity. In the Gi system, peak reductions in excitatory signaling were reached 30 min after injection and were sustained for at least 30 min thereafter (Figures S4B-S4D). In the Gq system, a 2.5 mg/kg (i.p.) dose of CNO increased excitatory neuronal calcium signaling by 42 ± 6.5%, while a 5.0 mg/kg dose increased signaling by 70 ± 8%. In the Gq system, there was a clear peak effect 30-45 min after injection (Figures S4B-S4D). In a control study, the highest dose of CNO used (5 mg/kg) did not alter neuronal calcium signaling in mice without DREADD transfection (Figure S4C). Based upon these tests, we adopted a 1.25 and 5.0 mg/kg dose of CNO in the Gi system, and a 2.5 and 5.0 mg/kg dose in the Gq system to impart modest or intermediate effects, respectively.

To this end, GCaMP6s;Cx3Cr1^CreER-eYFP^ mice were transfected with DREADD virus and microglial calcium activity was studied in the area of peak neuronal expression (mCherry tag; Figure 4A). In the Gi system, CNO-dependent decreases in CaMKIIa neuronal activity resulted in gradual increases in microglial process calcium activity, peaking 45-60 min after injection (Figures 4B-4D, and Video S4). Notably, there were clear dose-dependent differences in the extent of microglial process calcium activity after CNO injection (Figures 4C and 4D), with a 64 ± 11% increase observed at the lower dose (1.25 mg/kg) and a 133 ± 26% increase observed at the higher dose (5.0 mg/kg). In the Gq system, microglial process calcium activity displayed highly similar trends to the Gi system, despite the fact that these approaches resulted in opposing shifts to neuronal activity (hypoactivity in the Gi system and hyperactivity in the Gq system). At the lower dose in the Gq system, 2.5 mg/kg CNO administration resulted in a 68 ± 31% increase in microglial process calcium activity, while the higher dose (5.0 mg/kg) resulted in a 128 ± 13% increase in process calcium activity (Figures 4B-4D, and Video S4). In both the Gi and Gq systems, peak microglial calcium responses occurred 45-60 min after CNO administration, suggesting an approximate 15 min latency relative to peak neuronal changes (Figures 4D and S4D, gray highlighted regions). Additionally, both Gq- and Gi-based alterations to excitatory neuronal activity did not result in observable changes to microglial somatic calcium signaling (Figure 4D). Microglial process outgrowth was the predominant motility behavior observed following Gi- or Gq-DREADD activation in neurons, and extending processes had the greatest calcium activity (Figures 4E and 4F). Overall, we observe that a near-linear range of neuronal activity shifts (Figure S4C), results in a U-shaped curve of microglial process calcium activity (Figure 4C), with the greatest shifts in neuronal activity producing the largest microglial process calcium signaling.

**Figure 4:**
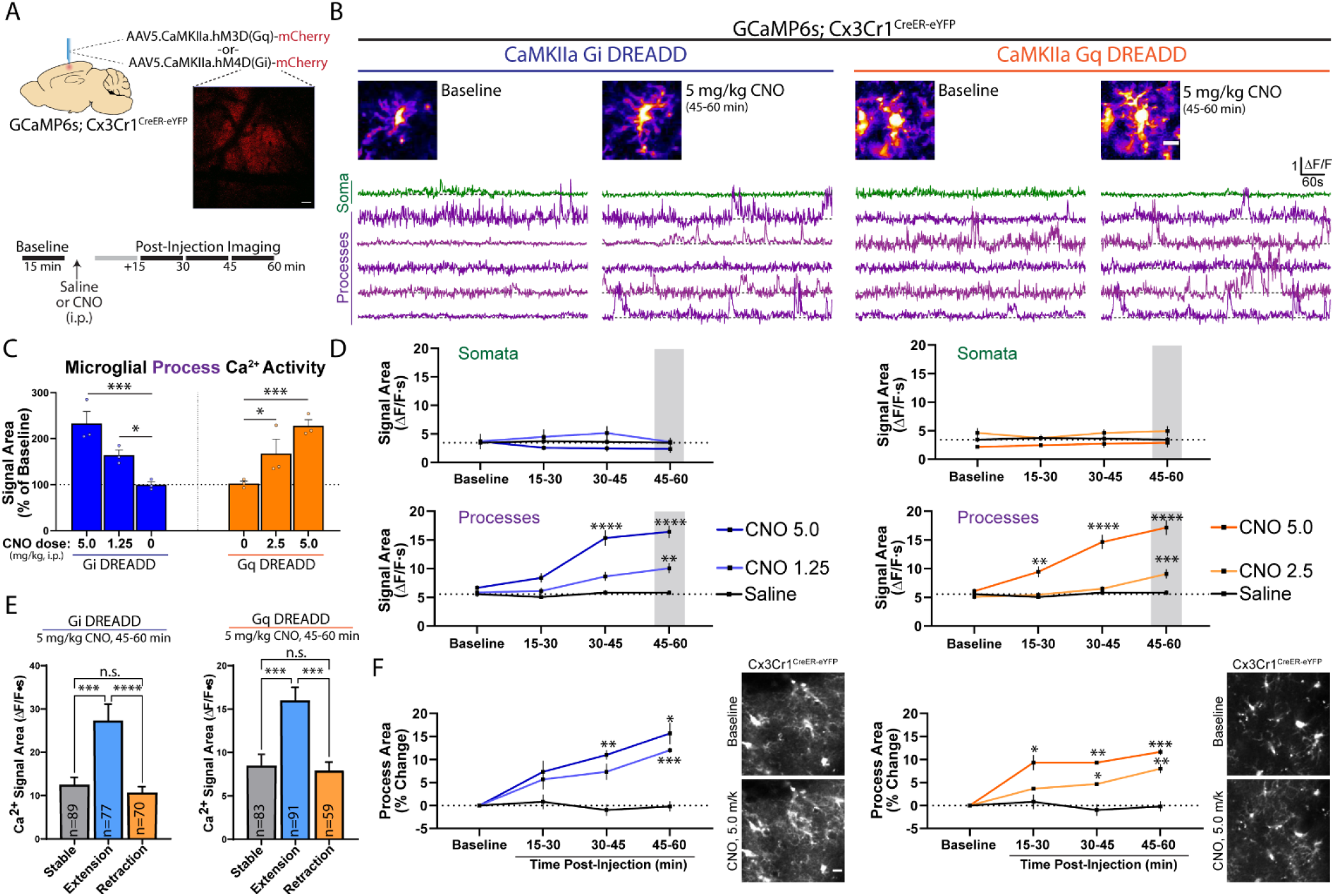
DREADD-based modulation of excitatory neuronal activity is sufficient to induce microglial calcium signaling. **(A)** Method to study neuronal DREADD effects on microglial calcium activity. mCherry expression served as a positive control for successful DREADD transfection. **(B)** Representative ΔF/F traces of microglial soma and process calcium activity at baseline and after CNO injection. **(C-D)** Saline or CNO-based effects on microdomain calcium activity, including (C) peak CNO-dependent changes in microglial process calcium activity (1-Way ANOVA with Dunnett’s post-hoc vs. saline), and (D) effects on both somata and processes over time (2-Way ANOVA with Dunnett’s post-hoc vs. baseline). **(E)** Microglial process calcium activity for processes undergoing extension, retraction, or remaining stable (1-Way ANOVA with Tukey’s post-hoc testing). **(F)** Changes in microglial process area and representative images of Cx3Cr1^CreER-eYFP^ morphology at baseline and following CNO administration (1-Way ANOVA with Dunnett’s post-hoc vs. baseline). Scale bars: 50 μm (A), 10 μm (B and F). N=6 GCaMP6s;Cx3Cr1^CreER-eYFP^ mice, half receiving Gq or Gi DREADD. Grouped data represent mean ± SEM; dots represent individual animals (C). Numbers of processes surveyed in (E) are shown in the bars. *p<0.05, **p<0.01, ***p<0.001, ****p<0.0001. See also Figure S4 and Video S4.

**Figure 5:**
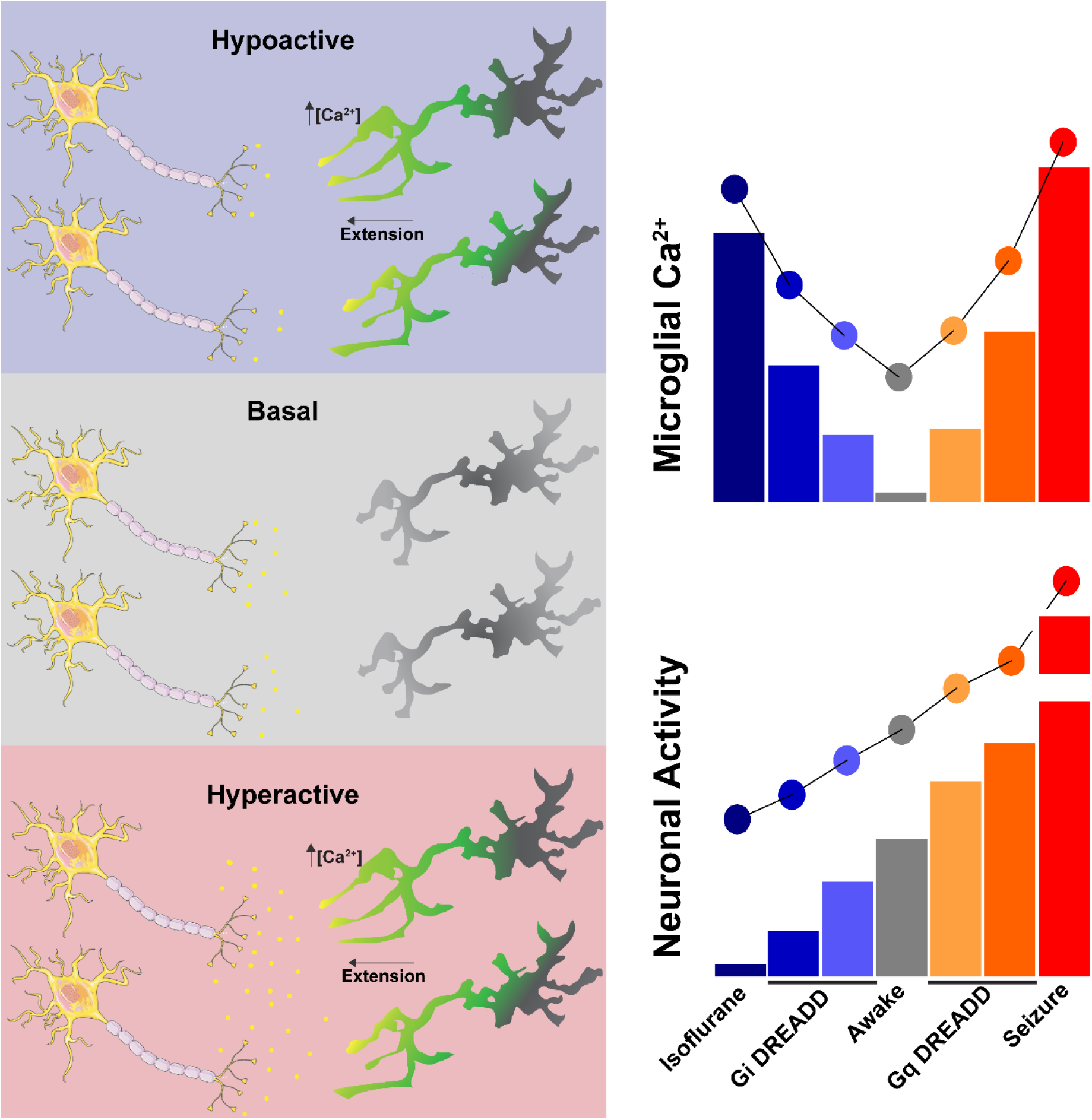
Summary of findings. **(Left)** Microglia display process extension and elevations of process calcium activity during neuronal hypoactivity or hyperactivity relative to their awake baseline. **(Right)** Microglia display a U-shaped curve of process calcium activity in relation to near-linear changes in neuronal activity.

## DISCUSSION

Herein, we describe a series of observations regarding microglial calcium signaling in response to shifts in neuronal activity. In general, microglia are considered to have the lowest levels of spontaneous calcium activity in “resting” or “baseline” states from reports in anesthetized mice (Brawek et al., 2017; Eichhoff et al., 2011; Pozner et al., 2015). We observe that microglial calcium activity is similarly low in awake, chronic window mice. However, pharmacological alterations in neuronal activity can trigger microglial process calcium signaling without impacting somatic calcium signaling. In general, microglial process calcium signaling follows a U-shaped curve, with greater shifts in neuronal activity (both hypoactive or hyperactive) resulting in stronger process calcium signaling. Additionally, both shifts in network activity were associated with microglial process extension. Extending processes consistently demonstrated the greatest calcium activity. This may explain why microglial calcium activity is minimal in the awake animal, as microglial processes are dynamically constrained in this state (Liu et al., 2019; Stowell et al., 2019).

Our findings clarify the role of neuronal activity on microglial calcium signaling. Previous studies using the GABA antagonist bicuculline to increase neuronal firing have come to different conclusions. In a first study, bicuculline did not alter microglial calcium activity (Eichhoff et al., 2011). However, this earlier study investigated bicuculline effects on microglial somatic calcium activity, deftly using a calcium indicator dye. Our work suggests that it is unlikely that acute changes in neuronal activity, even under seizure conditions, will dramatically impact microglial somatic calcium activity. In a more recent study using the calcium indicator GCaMP5G, microglia exhibited strong calcium wave activity when bicuculline was applied (Pozner et al., 2015). Notably though, changes in neuronal activity were evoked in an inflammatory context, specifically occurring 12-24h after LPS application, which suggests a synergy between inflammation and excitation on microglial calcium activity. We hypothesize that this observation is informative for interpreting our longitudinal studies following kainate status epilepticus. The period following a brain insult like status epilepticus alters neuronal network synchronization, but it can also provoke inflammation (Bar-Klein et al., 2014; Kim et al., 2017; Tian et al., 2017; van Vliet et al., 2018). Under these inflammatory and epileptogenic conditions, we similarly observe wave-like calcium activity in approximately 10% of imaging sessions, similar to the frequency described with bicuculline and LPS (Pozner et al., 2015). Notably, only under these long-term, pro-inflammatory conditions do we observe clear changes in microglial somatic calcium activity. The profoundly altered calcium signaling properties of microglia following status epilepticus suggests that calcium may serve as a key secondary messenger system in microglia during disease development.

The mechanisms regulating microglial calcium signaling have not been clearly identified. ATP is the most definitive agonist of microglial calcium signaling to date, and purinergic receptors (ionotropic P2X and metabotropic P2Y family) have been the most clearly implicated sensors for transducing microglial calcium signaling (Eichhoff et al., 2011; Färber and Kettenmann, 2006; Pozner et al., 2015). While kainate and the Gq DREADD system could reasonably increase the release of purinergic signaling molecules (Engel et al., 2016), evoking both process outgrowth (canonically P2Y12-driven, Eyo et al., 2014; Haynes et al., 2006; Swiatkowski et al., 2016) and process calcium activity (potentially P2Y6-driven, Koizumi et al., 2007), it would be illogical for Gi DREADD and isoflurane to increase purinergic signaling. Alongside Stowell et al., our recent studies suggest that depressing neuronal activity in the awake mouse prevents noradrenergic signaling to microglial β2 receptors (Liu et al., 2019; Stowell et al., 2019). Under awake physiology, β2 activation in microglia restrains process surveillance, motility, and extension, keeping microglia in a relatively arrested state. The absence of this signaling axis when neuronal activity is reduced allows for extension, as we observe in isoflurane and Gi DREADD studies. What is unclear, however, is whether β2 disinhibition also contributes to microglial calcium signaling during neuronal hypoactivity. Overall, our work demonstrates that microglia display increases in process calcium signaling in response to bidirectional changes in neuronal firing, suggesting that microglial calcium signaling is attuned to changes in neuronal activity states.

## ACKNOWLEDGEMENTS

We thank Dr. Vanda Lennon for early insights on project direction, and Dr. Greg Worrell and Yoga Varatharajah for assistance with code implementation. A.D.U. is supported by an NIH F32 grant (NS114040). L-J.W. is supported by the Mayo Foundation and NIH RO1 grants (NS112144 and NS088627).

## AUTHOR CONTRIBUTIONS

A.D.U. and Y.U.L. designed experiments. A.D.U. and L.L.B. performed experiments. A.D.U., L.L.B., and Y.Y. conducted data analysis. A.D.U. wrote the manuscript. L-J.W. supervised all phases of the project and provided funding.

## METHODS

### LEAD CONTACT AND MATERIALS AVAILABILITY

Further information and requests for resources and reagents should be directed to and will be fulfilled by the Lead Contact, Long-Jun Wu.

### EXPERIMENTAL MODEL AND SUBJECT DETAILS

GCaMP6s (Rosa26-CAG-LSL-GCaMP6s; 024106), Lck-GCaMP6f (Rosa26-CAG-LSL-Lck-GCaMP6f; 029626), and CX3CR1^CreER-eYFP^ (021160) knock-in mouse lines can be obtained from the Jackson Laboratory. Neuronal calcium activity was studied in C57BL/6J WT mice using AAV injections. Both male and female offspring were used across all studies at an age ranging from 3-5 months. Mice were group housed in an AAALAC-approved facility in climate-controlled rooms with a 12-h light/dark cycle (lights on at 6am) and had free access to food and water. All experimental procedures were approved by the Mayo Clinic’s Institutional Animal Care and Use Committee (IACUC).

### METHOD DETAILS

#### Administration of tamoxifen in chow

In CreER lines, tamoxifen was administered through chow to activate GCaMP expression in microglia. Mice were weaned at P21 and then provided tamoxifen in chow for a two week period (250 mg tamoxifen per 1 kg of chow; Envigo). Notably, chronic window studies occurred no sooner than 4 weeks after tamoxifen administration ended, which is sufficient to label microglia but not peripheral CX3CR1 cell types due to differences in their turnover rates (Parkhurst et al., 2013).

#### Stereotaxic delivery of AAV

Under isoflurane anesthesia (4% induction, 1.5-3% maintenance), AAVs were injected into the somatosensory cortex (AP: −4.5, ML: +2.0) using a glass pipette and micropump (World Precision Instruments). AAVs were targeted to both layer V neurons (DV: −0.5) and layer II/III neurons (DV: −0.2). A 250 nL volume was dispensed at each level at a 40 nL/min rate followed by a 10 min rest period for diffusion. pENN.AAV9.CaMKII.GCaMP6s.WPRE.SV40, AAV5.CaMKIIa.hM3D(Gq)-mCherry (1:1 ratio), and AAV5.CaMKIIa.hM4D(Gi)-mCherry were either injected alone (1:1 dilution in PBS) or in conjunction (1:1 ratio of DREADD and GCaMP virus). All viruses were used at a titer of 10^12^ GC/mL and acquired from Addgene.

#### Cranial window surgery

Cranial windows were implanted following standard techniques. Under isoflurane anesthesia (4% induction, 1.5-2.5% maintenance), a circular craniotomy (<3 mm diameter) was made over somatosensory cortex (AP: −4.5, ML: +2.0) using a high-speed dental drill. A circular glass coverslip (3 mm diameter, Warner) was secured over the craniotomy using Loctite 401 at the lateral edges. A 4-point headbar (NeuroTar) was secured over the window using dental cement. A mild NSAID solution (0.2 mg/mL Ibuprofen) was provided in the home cage for 72 h following surgery. In mice undergoing AAV injections, windows were implanted over the injection site after a 3-week recovery period.

### TWO-PHOTON IMAGING

#### Acute and chronic imaging parameters

Imaging in the awake animal was performed using a Scientifica multiphoton microscope equipped with galvanometer scanning mirrors. To image GCaMP fluorescence, a Mai-Tai DeepSee laser was tuned to 920 nm and kept below 10 mW output for layer I imaging or 12 mW output for layer II/III imaging. Imaging occurred at a 1 Hz frame rate at 512 x 512 pixel resolution using a 16x water-immersion objective (NA: 0.8, Nikon). Microglial calcium was imaged at a consistent zoom factor (300 x 300 μm area), while neuronal calcium was sampled from a larger area (450 x 450 μm).

#### Training and baseline imaging

Mice were trained to move on an air-lifted platform (NeuroTar) while head-fixed under a 2P objective. Training occurred for 30 min/day for the first 3 days following surgery and once per week thereafter. On the first day following surgery, a subset of mice was imaged to determine baseline microglial calcium activity in the acute window (Figure 1). Chronic imaging occurred no sooner than 4 weeks after surgery in mice with a clear window. Across all studies, mice were allowed 10 min to acclimate after being placed in the head restraint before imaging began. For studies of spontaneous calcium activity in awake acute and chronic window animals (Figures 1 and S1), a 15 min video was acquired 55-75 μm below the dura (layer I) and 150-170 μm below the dura (layer II/III).

#### Isoflurane studies

A cohort of chronic window GCaMP6s;Cx3Cr1^CreER^ mice and AAV9.CaMKII.GCaMP6s WT mice was first imaged under awake, baseline conditions (10 min; 55-75 μm depth for microglia, 150-170 μm depth for neurons; Figure 2). A nose cone was then secured to the head-restraint system and the mouse was induced with isoflurane on the platform (4%, up to 60 s). Isoflurane was maintained at 1.5-2% for the duration of imaging, acquired as three 10-min blocks. Under isoflurane, body temperature was maintained at 37°C (Physitemp).

#### Kainate studies

Two separate cohorts of GCaMP6s;Cx3Cr1^CreER^ mice were used to study either the acute effects of kainate during status epilepticus (Figure S2) or the longitudinal effects of KA-SE over a 14 d period (Figure 3). To study the acute effects of kainate status epilepticus, chronic window mice were imaged at baseline (15 min, 60-75 μm depth) then removed from the stage and injected with 19 mg/kg kainate (i.p.). Mice were then visually monitored for a generalized seizure (defined by Racine stage 3-5 criteria; Racine, 1972). The same region was then imaged for 30 min following the first generalized seizure, which began 45-75 min after injection. To study the longitudinal effects of status epilepticus, a 15 min baseline recording was taken from two separate regions of the cranial window (55-80 μm depth). Kainate was then administered (19 mg/kg, i.p.) and seizures were visually monitored. After 1 hr, booster injections (7.5 mg/kg, i.p.) were administered every 30 min to animals not displaying a generalized seizure until such time as mice displayed at least 8 generalized seizures. Chronic window mice meeting these inclusion criteria were then re-imaged in the same two locations 1, 2, 3, 7, 10, and 14 days after status epilepticus.

#### Chemogenetic studies

To validate the DREADD approach and determine CNO dose responses, we used WT C57BL/6J mice co-injected with AAV9.CaMKII.GCaMP6s and either AAV5.CaMKIIa.Gi-mCherry, or AAV5.CaMKIIa.Gq-mCherry. A cohort of mice injected with AAV9.CaMKII.GCaMP6s alone was employed for a CNO control study. To study the effects of CaMKIIa-DREADD effects on microglial calcium, we injected the Gq or Gi virus into GCaMP6s;Cx3Cr1^CreER^ mice. All experiments were performed in chronic window animals (minimum of 4 weeks after window surgery and 7 weeks after virus injection). Somatic changes in neuronal calcium were studied in layer II/III (150-170 μm; Figure S4), while microglia were studied in layer I (55-75 μm depth; Figure 4). As a positive control for microglial studies, imaging was only performed in a region with high mCherry expression (visualized with a 740 nm excitation wavelength), and this region remained consistent across all trials. A trial consisted of a 15 min acclimation period to head restraint, 15 min baseline imaging session, removal and injection (saline or 1.25, 2.5, or 5.0 mg/kg CNO, i.p.), 15 min re-acclimation period under head restraint, then three 15 min post-injection imaging blocks (see Figure 4A). Post-injection changes in calcium activity were compared to the daily baseline period. CNO trials were separated by at least 48 h, with lower doses administered first.

### IMAGE PROCESSING AND STATISTICAL ANALYSIS

#### Image processing, ROI selection, and calcium analyses

All videos were corrected in ImageJ using the template matching plugin with subpixel registration. An average intensity image was then generated for ROI selection. ROIs were manually drawn for all neuronal somata detected in layer II/III, which appeared as a light ring with a dark circular nucleus. Microglial somata were carefully outlined by hand. Microglial processes were either segmented by hand (Figures 1 and S1) or delineated with ImageJ thresholding (all other figures). In the latter case, microglial somata were first masked in the average intensity image, then a uniform threshold was applied to find the area of the processes. These ROIs are termed “microglial process territories” as they do not always represent a single process. Notably, the low-intensity but constitutive eYFP signal of microglia is preferentially captured in the average intensity projection, providing an image of cell morphology that is not strongly influenced by overall activity. Excel was used to convert fluorescent values into ΔF/F values and determine maximum amplitude and signal area. Baseline fluorescence was determined as the lower 25^th^ quartile value from a 100s moving window calculation. In some cases (microglial spreading calcium waves and neuronal calcium during kainate status epilepticus), the calcium transient could occur for a longer period than the moving window, so the trace was normalized to the 25^th^ quartile value of a 200-frame segment without major calcium activity. A calcium transient was considered to occur at a ΔF/F threshold over three times greater than the standard deviation of the baseline. The signal area represents the sum of any ΔF/F value above this threshold across all frames. A soma or process was considered active if it had a signal area of ≥5 ΔF/F·s.

#### Motility and Sholl analyses

In short-term experiments, morphology changes were determined by finding the overall process area for each time period (Figures 2, 4, and S2). Using average intensity projections of the eYFP signal (full 600 or 900 frames), microglial somata were masked and a uniform threshold was applied to the determine process area. In longitudinal or long-term experiments (Figures S1 and S3), morphological complexity was determined using the Sholl analysis plugin of ImageJ (Ferreira et al., 2014). To determine process motility behavior, a 30-frame moving window average was compiled from each T-series. Clearly identifiable microglia (soma and processes in the field of view) were then selected from the first frame (n=5-10 cells/animal). The motility of each primary and secondary process was first visually investigated, and an ROI was manually created around the maximum area of the process (>9 μm^2^ size inclusion criteria). Motility behavior was then definitively assigned by applying a uniform threshold and obtaining area values for each process ROI. Processes undergoing extension were characterized by a >50% increase in area, while processes undergoing retraction were characterized by a >50% decrease in area. Stable processes exhibited less than a 50% change in area in either direction. Dynamic processes undergoing both a >50% increase and decrease in area were excluded. The associated calcium activity for each manually segmented process was determined from the full 1Hz T-series video.

#### General statistical analysis

Prism (GraphPad, version 8) was used for group analyses. In most experiments, statistical significance was determined using a one- or two-way ANOVA design with Dunnett post-hoc comparison to the values obtained in the baseline period. Differences in the proportion of active microdomains were assessed using a Fisher’s exact test. All data are expressed as the mean ± SEM. Statistical details are provided in the figures and figure legends.

