## Supplementary material for "Microglial Calcium Signaling is Attuned to Neuronal Activity": Figure S

**Supplemental Material**

**Supplemental Figures**

**Figure S1. Related to Figure 1. Spontaneous microglial calcium activity differs between acute and chronic window preparations. (A)** Timeline of acute and chronic window imaging and animal training.

**(B)** eYFP average intensity images of microglia morphology in acute and chronic window preparations. Sholl analysis of microglial morphology (n=50 microglia per group; 2-Way ANOVA).

**(C-D)** Microglial ∆F/F calcium traces from a representative microglial cell in an acute window animal (C) or cells in a chronic window animal (D).

**(E)** Percent of active microdomains (Fisher’s exact test).

**(F)** The amplitude recorded from microdomains (1-Way ANOVA with Sidak’s post-hoc comparison).

**(G)** The calcium signal area recorded from microdomains (1-Way ANOVA with Sidak’s post-hoc comparison). Cumulative distribution curves for Layer I and Layer II/III process calcium signal areas.

**(H)** Microglia soma and process calcium signal areas recorded over three successive days in chronic window animals

Scale bars: 50 µm (B). Grouped data represent the mean ± SEM; dots represent microdomains (F and G); dashed lines represent individual animals (H). The number of animals and microdomains surveyed is described in (E). **p<0.01, ***p<0.001, ****p<0.0001.

**Figure S2. Related to Figure 3. Increased microglial process calcium activity immediately following the observation of kainate-induced seizures.**

**(A)** Outline of the experimental timeline.

**(B-C)** CaMKIIa excitatory neuronal calcium activity (AAV.CaMKIIa.GCaMP6s.WPRE transfection) in the somatosensory cortex before and after kainate administration. (B) Representative ∆F/F traces at baseline, 15-30 min following kainate, and after the first observed generalized seizure. (C) Signal areas derived from CaMKIIa somata in layer II/III (1-Way ANOVA with Dunnett’s post-hoc comparison to baseline).

**(D-E)** Microglial calcium activity (GCaMP6s;Cx3Cr1^CreER-eYFP^) in the awake animal at baseline and after the first kainate-induced generalized seizure. (D) Representative ∆F/F traces from microglial somata and processes and the percentage of active microdomains under each condition (Fisher’s exact test). (E) Summary of calcium signal area values at baseline and after the first generalized seizure (2-Way ANOVA with post-hoc comparison between kainate and baseline).

**(F)** Cx3Cr1^CreER-eYFP^ average intensity images of microglial morphology at baseline and after kainate-induced seizures. Changes in process area between conditions (paired t-test).

Scale bars: 50 µm (B), 10 µm (D). Grouped data represent the mean ± SEM; dots represent individual neuronal somata (C) or individual microdomains (E); dashed lines represent individual animals (F). N=3 mice for neuronal calcium studies (B, C), N=4 mice for microglial calcium studies (D-F); the number of microdomains surveyed is provided in (D). *p<0.05, ****p<0.0001.

**Figure S3. Related to Figure 3. Longitudinal changes in microglial morphology following kainate status epilepticus.**

**(A)** Sholl analysis profiles of microglial morphology 1, 2, 3, 7, 10, and 14 days following kainate status epilepticus plotted against the baseline period (2-Way ANOVA, main effect of KA).

**(B)** Summary of longitudinal changes in morphology created by summing the total number of branch point intersections determined by the Sholl analyses in (A).

Data represent the mean ± SEM. 50 microglia surveyed from N=5 mice at each time point. *p<0.05, ****p<0.0001; p values represent the main effect of KA on morphology from 2-Way ANOVA analyses of Sholl distributions for each time period studied.

**Figure S4. Related to Figure 4. Characterization of DREADD-based changes in CaMKIIa neuronal calcium activity.**

**(A)** AAVs injected into the somatosensory cortex of WT mice in order to study DREADD-based changes in neuronal calcium activity (top). Experiment outline (bottom).

**(B)** Representative ∆F/F traces of CaMKIIa neuronal calcium activity at baseline and after CNO injection.

**(C-D)** Saline or CNO-based effects on neuronal calcium activity, including (C) peak CNO-dependent changes in CaMKIIa calcium activity in the Gi and Gq systems (1-Way ANOVA with Dunnett’s post-hoc comparison to saline), and (D) effects over time (2-Way ANOVA with Dunnett’s post-hoc comparison to baseline).

Scale bar: 50 µm (B). N=8 WT mice, N=4/8 receiving AAV-CaMKIIa-hM3D(Gi) and N=4/8 receiving AAV-CaMKIIa-hM4D(Gq). Grouped data represent mean ± SEM; dots represent individual animals (C). *p<0.05, **p<0.01, ***p<0.001.

**Supplemental Figure 1**

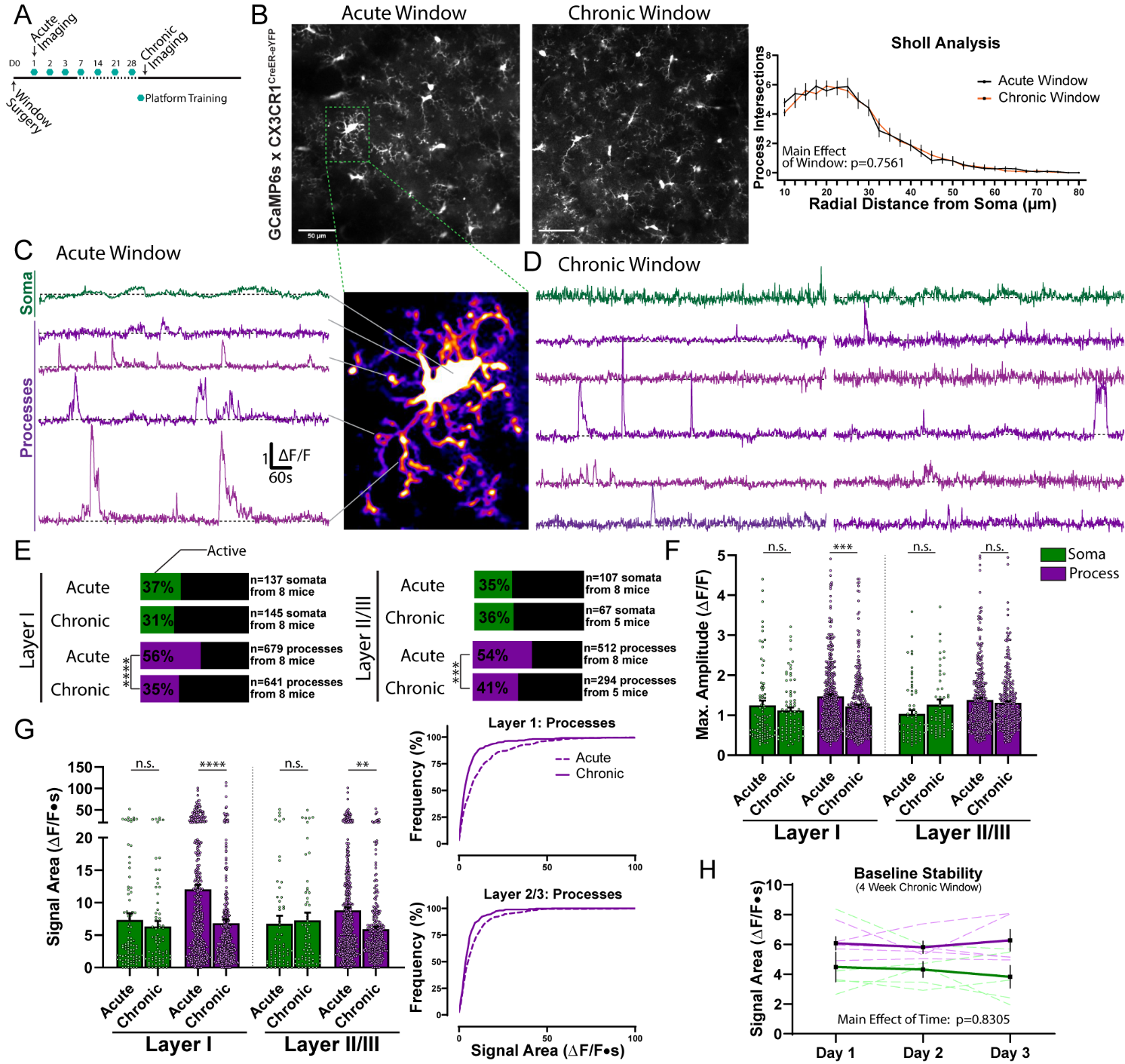

**Supplemental Figure 2**

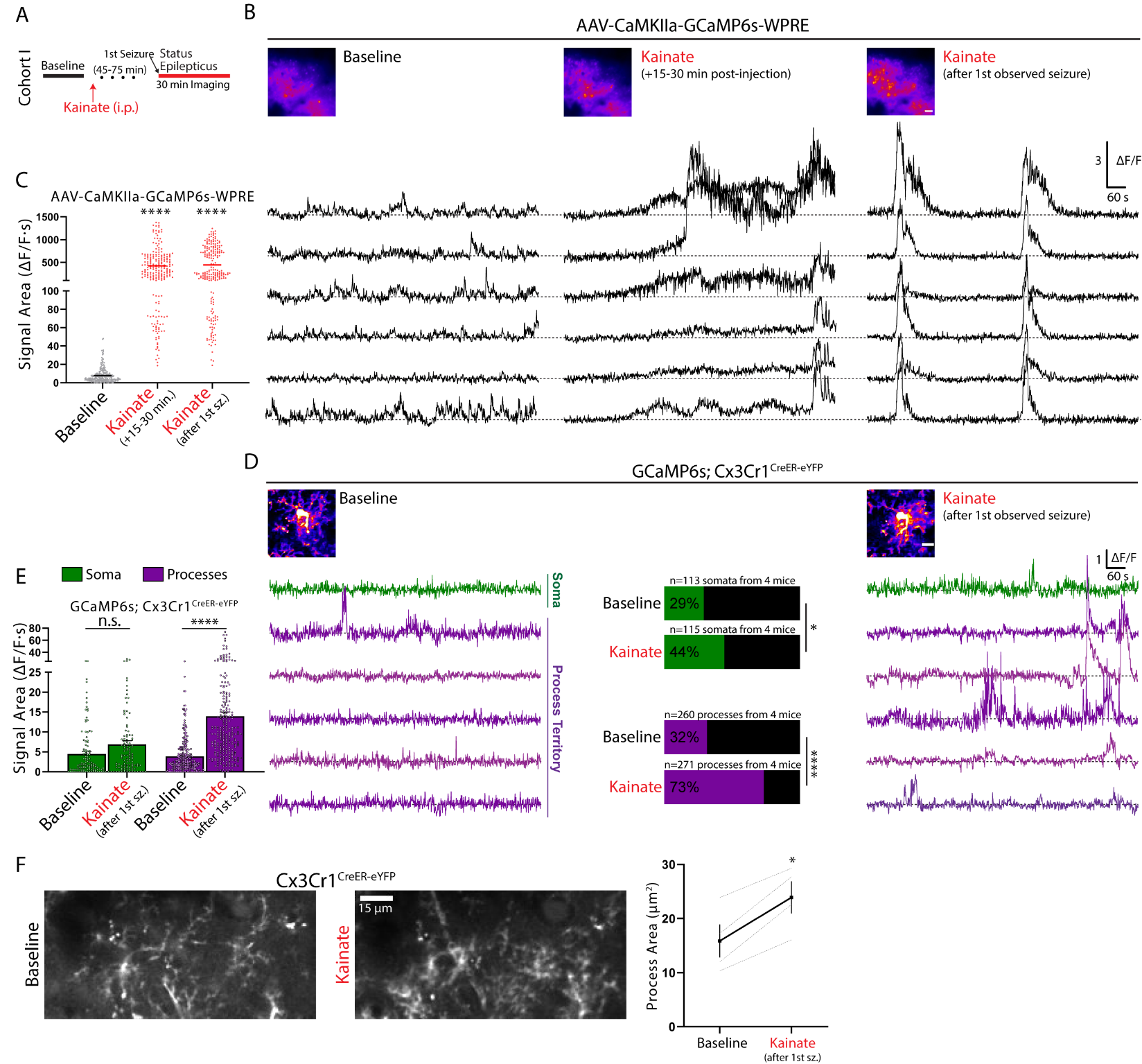

**Supplemental Figure 3**

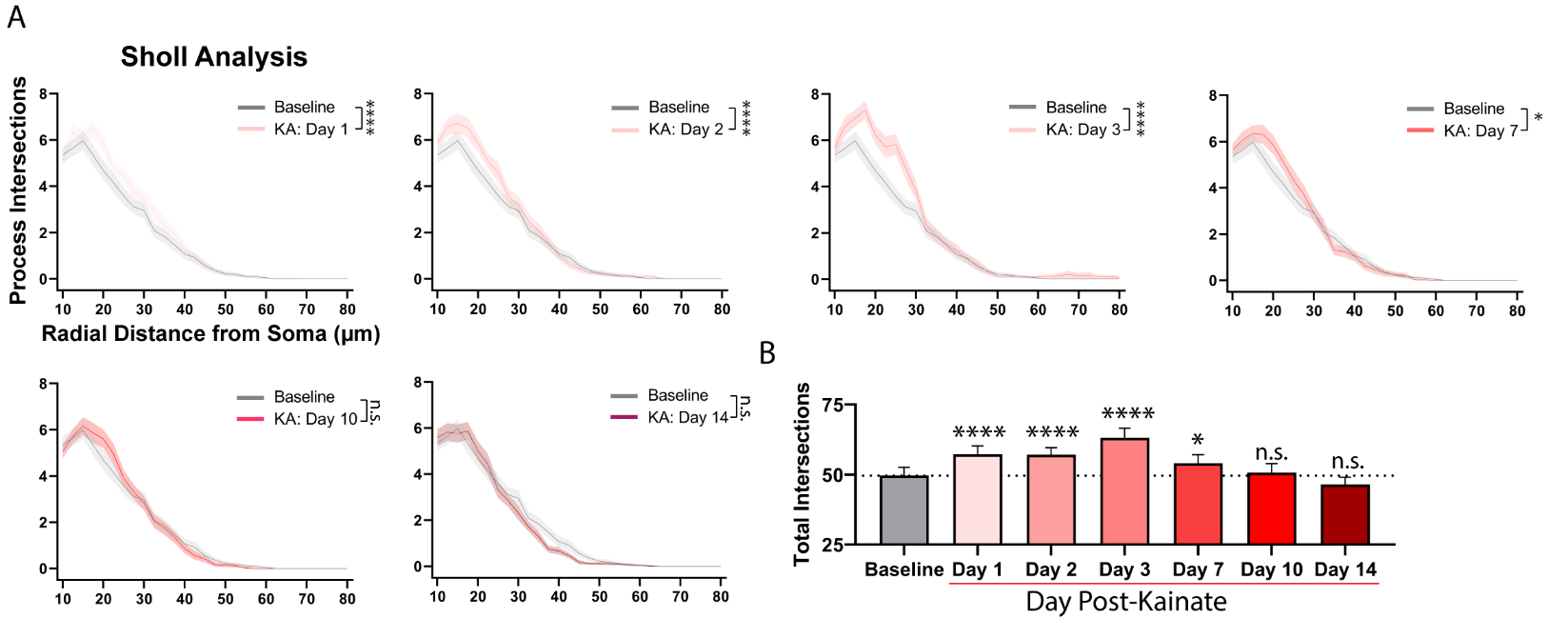

**Supplemental Figure 4**

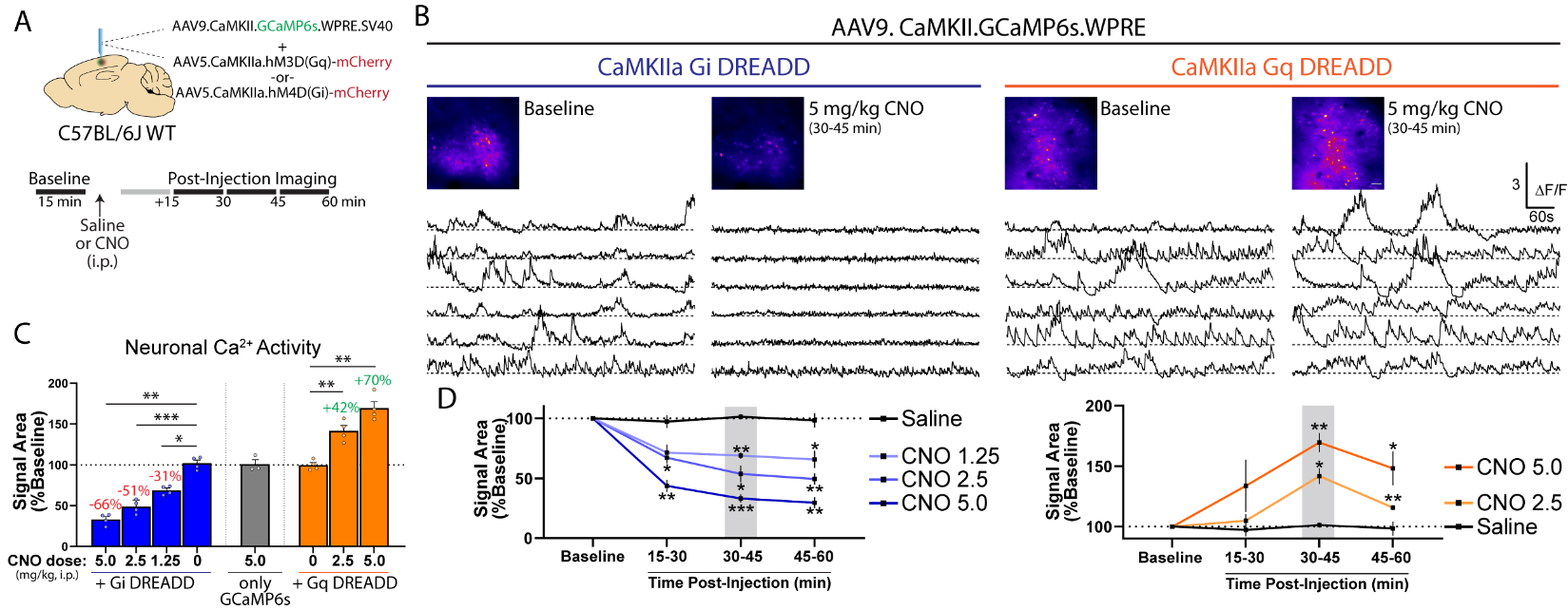

**Supplemental Videos**

**Video S1. Related to Figure 2. Microglial calcium changes in the awake mouse and during 30 min isoflurane administration.**

**Video S2. Related to Figure 3. Spreading microglial calcium waves recorded one day after kainate status epilepticus.**

**Video S3. Related to Figure 3. Microglial process extension co-occurring with increased calcium activity.**

**Video S4. Related to Figure 4. Microglial calcium activity at baseline and following CNO administration in Gi and Gq systems.**
